# An Expert-Curated Benchmark for Frontier and Local LLMs Querying a Biomedical Knowledge Graph

**DOI:** 10.64898/2026.10.04.756532

**Authors:** Johannes F. Medagbe, Bonface M. Kilyungi, Arun Isaac, Alexander Kabua, Felix Lisso, Sammy Wambua, Robert W. Williams, Pjotr Prins

## Abstract

We combine GeneNetwork’s 1TB structured database with its metadata as an RDF graph and an associated literature corpus. We benchmark frontier and local LLMs querying it through four architectures: document, graph and agentic retrieval-augmented generation (RAG), and a combination of the three. We evaluate them on 40 domain specific expert-curated questions with a new metric: the satisfaction frequency ***ψ*** (*psi*, i.e, the share of questions answered with precision and recall of at least 70% or an f1 of at least 60%). ***ψ*** ranks systems the same way precision, recall and f1 — a machine learning metric built from the harmonic mean of precision and recall — but is easier to read and assess for humans. System architecture had a large effect: with full use of Claude Opus 4.8 the plain document RAG of our earlier GNQA system had a ***ψ*** of 36% against 86% for our new combined tools. Delegating most of the work from Claude Opus 4.8 to a local model halved query costs while still reaching a ***ψ*** of 59%.

## 1 Introduction

GeneNetwork (GN)(Mulligan et al. 2017) is both a FAIR data repository, SQL database, and analytic platform for systems genetics, containing over 25 years of experimental genotype, phenotype and molecular trait data across mouse, rat, human and other species. GN offers a shared toolkit for QTL and eQTL mapping, correlation analysis and network reconstruction through a user-friendly interface (UI) (Watson and Ashbrook 2020; Parker et al. 2017; Mulligan et al. 2017; Lu and Lu 2013). The SQL database is internal and, despite exposing UI and APIs, data in GN is not always easy to navigate and digest, thereby not only limiting accessibility by researchers, but also discoverability by Large Language Models (LLMs). We aimed to address such limitations by modeling GN metadata as a graph in the form of the resource description framework (RDF) and make those available through a discoverable SPARQL REST API.

This work builds directly on our earlier published GeneNetwork Question-Answer system, GNQA (Darnell et al. 2024). GNQA introduced a GPT-driven, document-level RAG over a corpus of three thousand peer-reviewed publications on GN, aging and/or diabetes. GNQA demonstrated that returning provenance alongside every answer, i.e. the exact paragraphs and papers an answer was drawn from, meaningfully reduced hallucination and improved trust among both domain experts and citizen scientists (Darnell et al. 2024). GNQA had three notable limitations, however, that motivated the present study. First, GNQA only indexed unstructured text describing GN; it never gave the LLM direct access to GN’s own structured experimental data, so questions requiring the underlying dataset itself, rather than papers about it, remained out of reach. Second, GNQA relied on a single, naive document RAG architecture; domain experts who evaluated the system explicitly suggested that agent-based approaches could improve response quality, a suggestion we act on here. Third, GNQA was built on a closed, proprietary stack (OpenAI’s GPT and FahamuAI’s RAG-as-a-service), which the original authors flagged as a limitation to be addressed with open-source models and composable RAG engines in future work.

LLMs can reproduce complex reasoning and accelerate link discovery for biomedical research (Aamer et al. 2025; Luo et al. 2025; Zhou et al. 2025; Lu et al. 2024). A recently published effort, LongevityBench, benchmarked frontier and compact LLMs on their ability to reason from structured experimental data rather than text alone (Zhavoronkov et al. 2026). In our study, we combined RDF data, generated directly from GN’s metadata in the structured database, with its associated literature corpus to enrich context for LLMs because RDF syntax is very similar to natural language (Tomaszuk and Hyland-Wood 2020; Gerber and Ngomo 2014). Thus we created a valuable interface for data systems where one can move from RDF to natural texts and vice versa (Regino et al. 2026; Smeros et al. 2025; Rangel et al. 2024; Chen et al. 2021; Zou et al. 2014).

We designed and compared four systems over this RDF+literature resource: a document RAG, a graph RAG, an agentic graph RAG and a combination of all three (called ‘combi’ throughout the paper), going beyond GNQA’s single document RAG design. The document RAG follows the plain retrieve-then-generate design GNQA already used, fetching text passages by similarity and answering from them in a single pass. The graph RAG and the agentic graph RAG are cognitive architectures, meaning the model plans and acts on the structured data, writing SPARQL queries against the RDF graph and, in the agentic case, inspecting the results and revising its queries over several steps before it answers. The hybrid RAG runs the document, graph and agentic systems together and merges their answers (‘combi’), and it can route between a local and a frontier model so that the costly model is called only for the harder steps.

Finally, we benchmarked frontier and local models querying the database and literature corpus, to address GNQA’s call for evaluating open-source alternatives and reducing search cost. By frontier models we mean the large proprietary systems reached through commercial APIs, such as Anthropic’s Claude. By local models we mean the open-weight systems small enough to run on local GPUs (in this case a consumer grade NVIDIA RTX 3090 with 24Gb VRAM), such as Qwen3 or GPT-OSS. The frontier and local categories differ in raw capability, in cost per query and in how much control a research group keeps over its own inference, so we treat the choice of model as a comparison axis in its own right, alongside the system architecture. To compare the resulting systems we built a benchmark of 40 domain specific questions with reference answers, curated by domain experts from GN’s resource classification scheme. An LLM judge scores each generated answer against its reference for precision and recall, which we combine into a single metric, the *satisfaction frequency ψ*: the fraction of benchmark questions a system answers well enough to satisfy a user. We also record response time and token use, since a search tool has to be fast and affordable to be adopted. All tools and scripts are available online and published under the liberal BSD free software license.

## 2 Results

### 2.1 We introduced the performance metric *ψ* for humans

Performance evaluation of LLM systems typically calls for the use of specific metrics (accuracy, precision, recall, f1). In our setting an LLM judge compares each generated answer to its reference and counts the statements they share (true positives), the extra statements in the answer (false positives) and the reference statements it misses (false negatives). Precision is then TP*/*(TP + FP), recall is TP*/*(TP + FN) and the f1 score is their harmonic mean, while accuracy is not reported because true negatives cannot be counted (section 4.3.3). We believe that digesting four scores is not intuitive to the average reader. We therefore customized standard classification metrics into a new metric that informs on the number of questions a given system produced satisfactory answers for. We refer to the metric throughout the paper as the satisfaction frequency *ψ*. The reasoning is that a system with a higher satisfaction frequency *ψ* is more likely to meet users expectations.

To come up with a single metric we benchmarked experiments to compare evaluation results with the standard metrics (precision, recall, f1) using local models (Qwen3 30B A3B and GPT OSS 20B). Figure 1 shows performance results across LLM systems with the standard and customized metrics, respectively. Precision, recall and f1 reported that the combi system performed best and is followed by the document RAG. The same findings were confirmed with the satisfaction frequency *ψ*. In addition, both the standard and customized metrics advocated for the superiority of Qwen3 30B A3B over GPT OSS 20B on most systems with the exception of the graph RAG. Together, these patterns demonstrated that the customized metric provides similar insights to the standard metrics. Therefore we adopted it as the new standard for subsequent performance evaluation tasks.

**Fig. 1.**
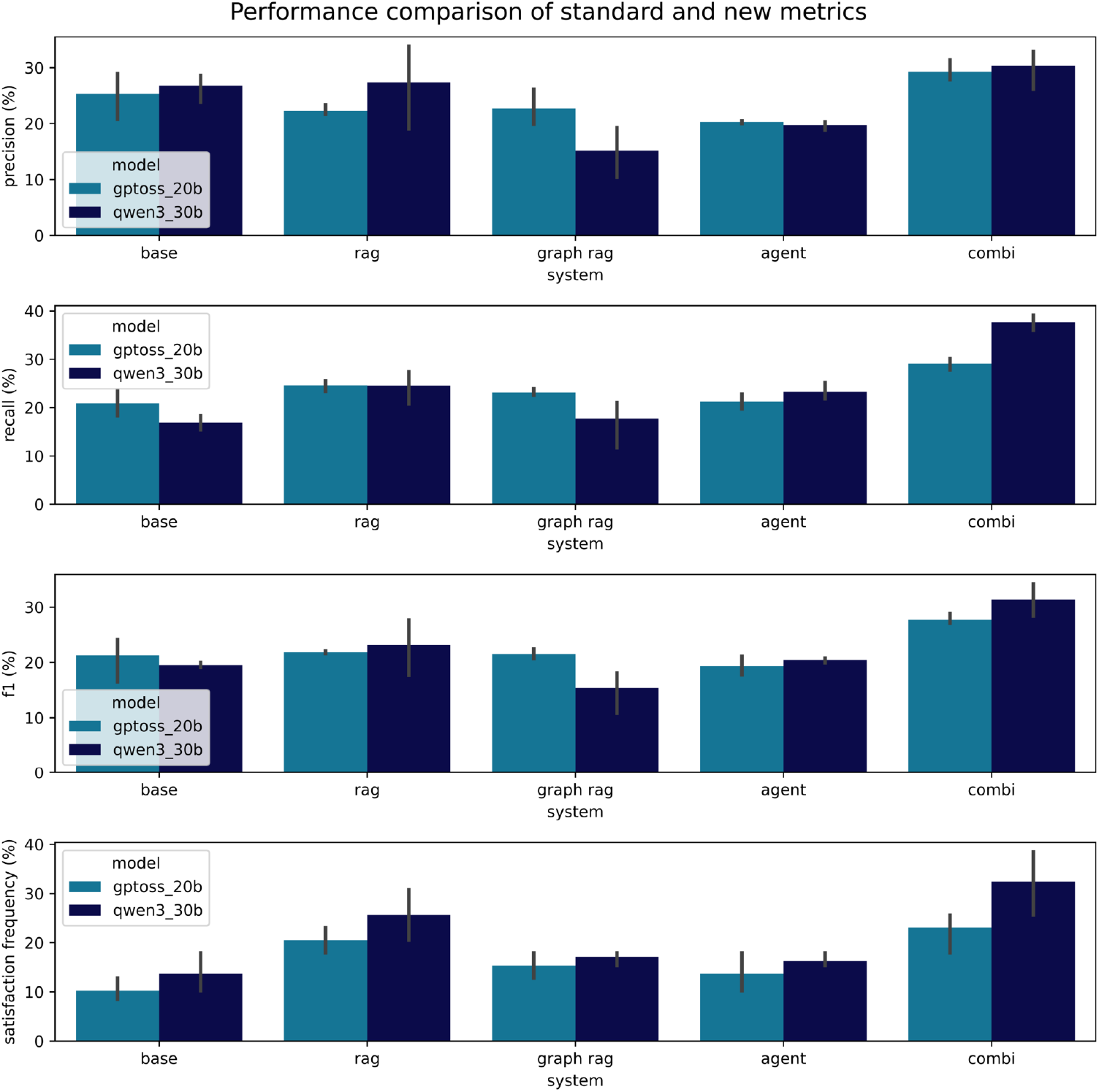
Precision, recall, f1 and satisfaction frequency *ψ* compared (top to bottom panels) for the bare language model (*base*) and the four GN LLM systems, document RAG (*rag*), graph RAG (*graph*), agentic graph RAG (*agent*) and combi RAG (*combi*), each run with the local models GPT-OSS 20B (steel blue) and Qwen3 30B A3B (midnight blue) on the 40-question expert benchmark. For every answer an LLM judge counted the statements shared with the reference (true positives), the extra statements (false positives) and the missing ones (false negatives) to compute precision, recall and f1 scores. Satisfaction frequency *ψ* is the percentage of benchmark questions a system answered well enough to count as satisfactory, defined as precision and recall of at least 70% or an f1 of at least 60%. Bars are medians over the 40 questions and three repeats and error bars show the spread. The combi system scores highest on all three metrics with either model. The document, graph and agentic RAG stay close to the bare model on these metrics. Qwen3 30B A3B leads on most systems, while GPT-OSS 20B is ahead on the graph RAG.

### 2.2 Architecture sets quality: the ‘combi’ leads and document RAG trails

We assessed the impact of the document RAG, graph RAG and agent architectures on feedback quality using the new metric validated in section 2.1. Each system was instantiated with a series of models (Qwen3 30B A3B, GPT-OSS 20B, Claude Haiku 4.5 and Claude Opus 4.8) and executed on questions in the benchmark over three iterations to provide a good baseline for architecture comparison.

All systems in figure 2 performed better than the base language model, demonstrating the value of building systems around LLMs. Overall, the architectures were ranked as follows: combi, agent, graph RAG and document RAG. LLM systems composing directly with the RDF knowledge base (graph and agentic RAG) substantially improved inference, especially when powered with a very large model such as Claude Opus 4.8.

**Fig. 2.**
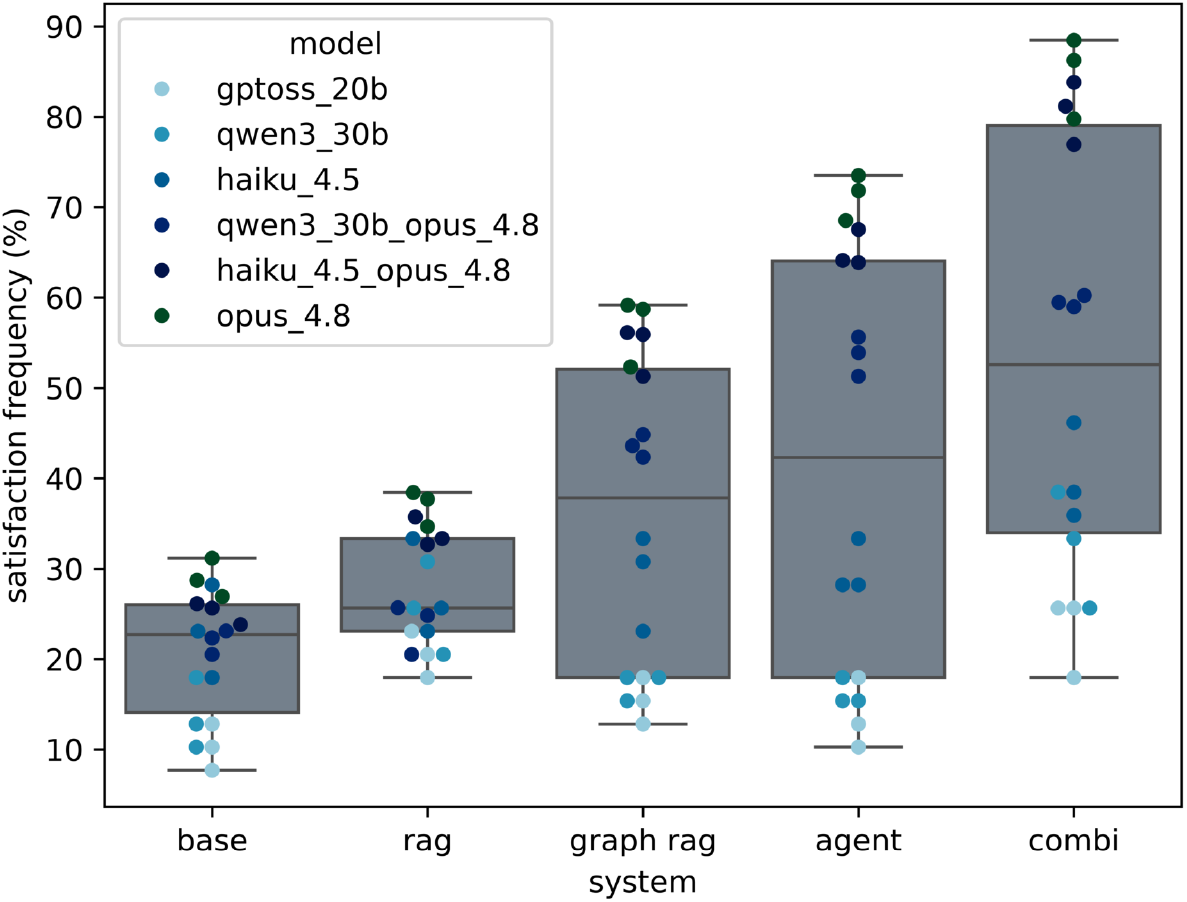
Satisfaction frequency *ψ* of the bare language model (*base*) and the four GN LLM systems, document RAG (*rag*), graph RAG (*graph*), agentic graph RAG (*agent*) and combi RAG (*combi*) on the 40-question expert benchmark; each box summarises satisfaction frequency of the architecture across the four models (Qwen3 30B A3B, GPT-OSS 20B, Claude Haiku 4.5 and Claude Opus 4.8) and three benchmark repeats. Satisfaction frequency *ψ* is the percentage of benchmark questions answered satisfactorily; boxes show the median and interquartile range over three repeats, whiskers the full range; each data point corresponds to a run with a specific model. Median satisfaction frequency *ψ* rises from the bare model through the document, graph and agentic RAG to the combi.

Our combined architecture (combi) exceeded in performance the agentic RAG for every model we tested. A paired t-test on the three repeats (section 4.3.2) confirmed this difference for Claude Opus 4.8 (+13.5%, *t*(2) = 11.6, *p* = 0.007) and Qwen3 30B A3B (+16.2%, *t*(2) = 5.3, *p* = 0.034), but not for Claude Haiku 4.5 (+10.3%, *t*(2) = 2.6, *p* = 0.12) or GPT-OSS 20B (+9.4%, *t*(2) = 2.0, *p* = 0.19); with only two degrees of freedom per test, the latter two should be read as inconclusive rather than as evidence of no real difference. This illustrates that despite the capabilities of cognitive architectures, simpler architectures also hold some merit. They can offer an extra boost of performance to agentic systems, in addition to lowering cost and addressing privacy concerns; such as signing up and submitting documents, data and queries to AI companies.

We identified an interaction between the LLM system architecture and the model used: the frontier-versus-local gap in satisfaction frequency *ψ* was small on the document RAG but large on the combi system. Averaging the two frontier models (Opus 4.8, Haiku 4.5) against the two local models (Qwen3 30B A3B, GPT-OSS 20B) within each repeat, this frontier-local gap was 6-15% on the document RAG versus 30-38% on combi; a paired t-test on the three repeats’ gap-of-gaps confirmed the interaction (*t*(2) = 4.8, *p* = 0.042). This explains the variability observed in figure 2 which corresponds to differences in satisfaction frequency *ψ* of the systems with large frontier and small local models. We zoomed in the interactions in figure 5 and provided an in-depth analysis in section 2.4.

### 2.3 The stronger architectures are slower, and latency does not track token use

We conducted additional experiments with the local models to gauge performance during deployment. Optimal search systems should provide state-of-art generation but also balance speed and token consumption. A system that gives very good answers but has a long generation time and high token consumption, might limit widespread adoption. From figure 3, it is clear that advanced systems take longer to run. The combi, agentic and graph RAG all required more work than the document RAG.

**Fig. 3.**
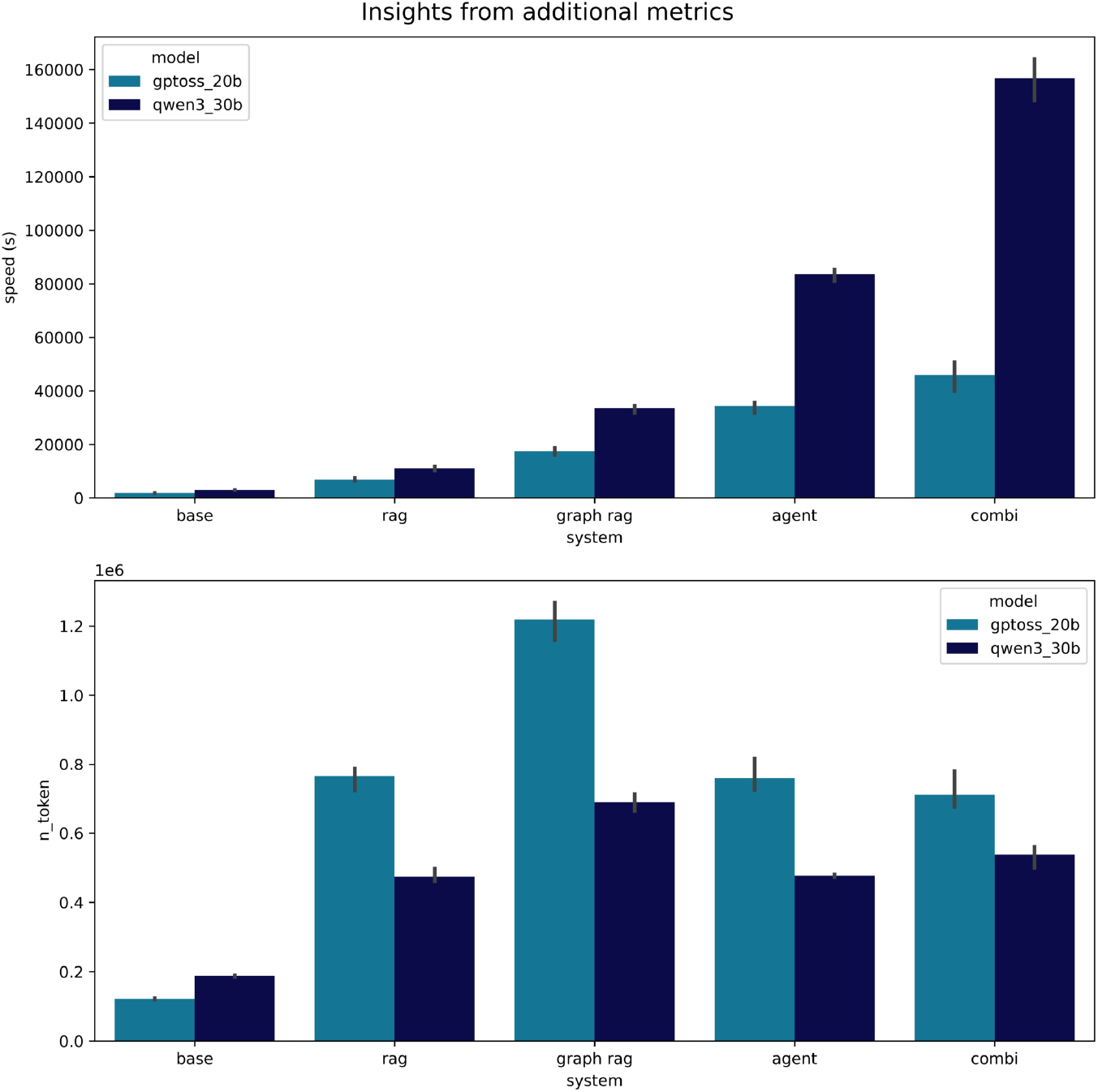
Wall-clock time per query (top, seconds — less is better) and total tokens consumed (bottom) for the bare language model (*base*) and the four GN LLM systems, document RAG (*rag*), graph RAG (*graph*), agentic graph RAG (*agent*) and combi RAG (*combi*), each run with the local models GPT-OSS 20B (steel blue) and Qwen3 30B A3B (midnight blue) on the 40-question benchmark; bars are medians over the 40 questions and three repeats and error bars show the spread. Query time grows steadily with architecture complexity, from a few seconds for the bare model to several minutes for combi, and Qwen3 30B A3B is markedly slower than GPT-OSS 20B on the agentic and combi systems. Token use does not follow the same order: the graph RAG consumes the most tokens yet runs faster than the agentic and combi systems, which reuse context and so spend fewer tokens despite taking longer.

Interestingly, the level of work in terms of processing time did not quite correlate with token usage. This was not dependent on the model used for generation. For instance, the agentic and combi systems consumed less tokens than the document and graph RAG. Based on those observations, we decided to address the latency of combi and motivate its adoption by the community. We introduced output streaming to cut down time. Figure 4 depicts streaming in each system during query handling.

**Fig. 4.**
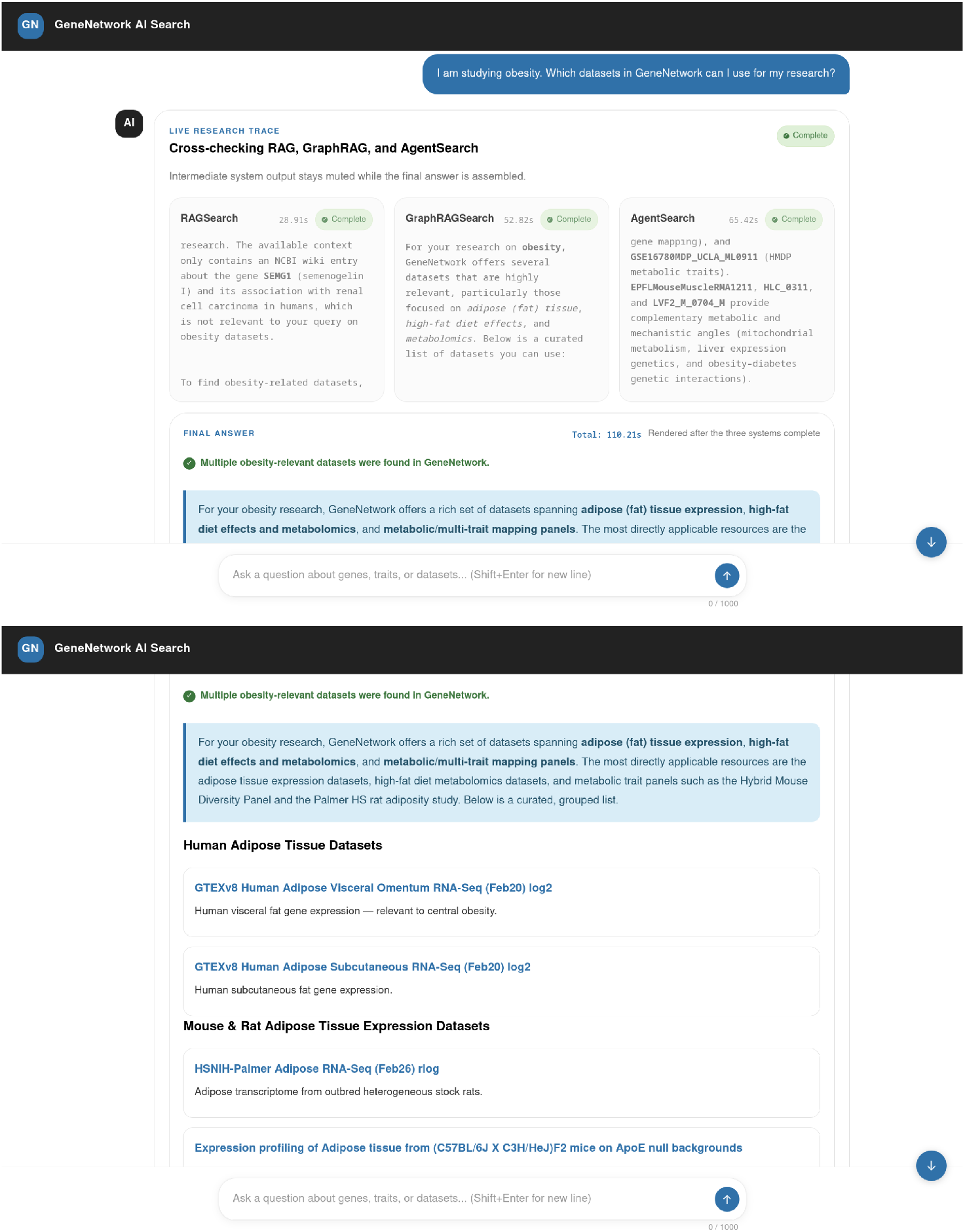
Screenshots of the GN AI Search web interface answering the query *I am studying obesity. Which datasets in GeneNetwork can I use for my research?* Combi runs by default: the document (RAGSearch), graph (GraphRAGSearch) and agentic (AgentSearch) subsystems run in parallel, each streaming its intermediate output with its own completion time (here 28.9, 52.8 and 65.4 seconds), and a final answer is synthesised once all three finish (110.2 seconds total). The lower panel shows the assembled answer, a grouped and annotated list of relevant GN datasets. Earlier turns are kept in context so follow-up questions can build on them.

### 2.4 Frontier models outperform local ones, but a local-frontier pair recovers most of the gap at lower cost

The performance of any LLM system heavily relies on the capabilities of the model that it leverages. In this work, we aimed to build sustainable LLM systems for biomedical research with GN. For this reason, we preferred experimenting with local models over the numerous frontier models that exist. We limited our selection of local models to Qwen3 30B A3B and GPT-OSS 20B because of GPU limitations (24Gb VRAM). We compared those models with a small frontier model (Claude Haiku 4.5). To provide a point of reference with larger frontier models, we also evaluated performance with Claude Opus 4.8.

Our findings confirmed the big gap between the local models we selected (Qwen3 30B A3B and GPT-OSS 20B) and Anthropic models (see figure 5). Claude Haiku 4.5 without our systems showcased a better performance than both Qwen3 30B A3B and GPT-OSS 20B. Yet, they have comparable sizes. Model size alone cannot explain this, but differences in architecture, tokenization strategy, pretraining data and Reinforcement Learning from Human Feedback (RLHF) regime across model families could; also the local models were run at reduced precision (Q4 K M and F16) rather than full precision, so we cannot isolate precision impact. Qwen3 30B A3B outperformed GPT-OSS 20B across all systems in figure 5. This was also observed in figure 1. We attributed disparities between the two models to the number of parameters mostly. However, we do not think the relationship between performance and number of parameters is linear.

**Fig. 5.**
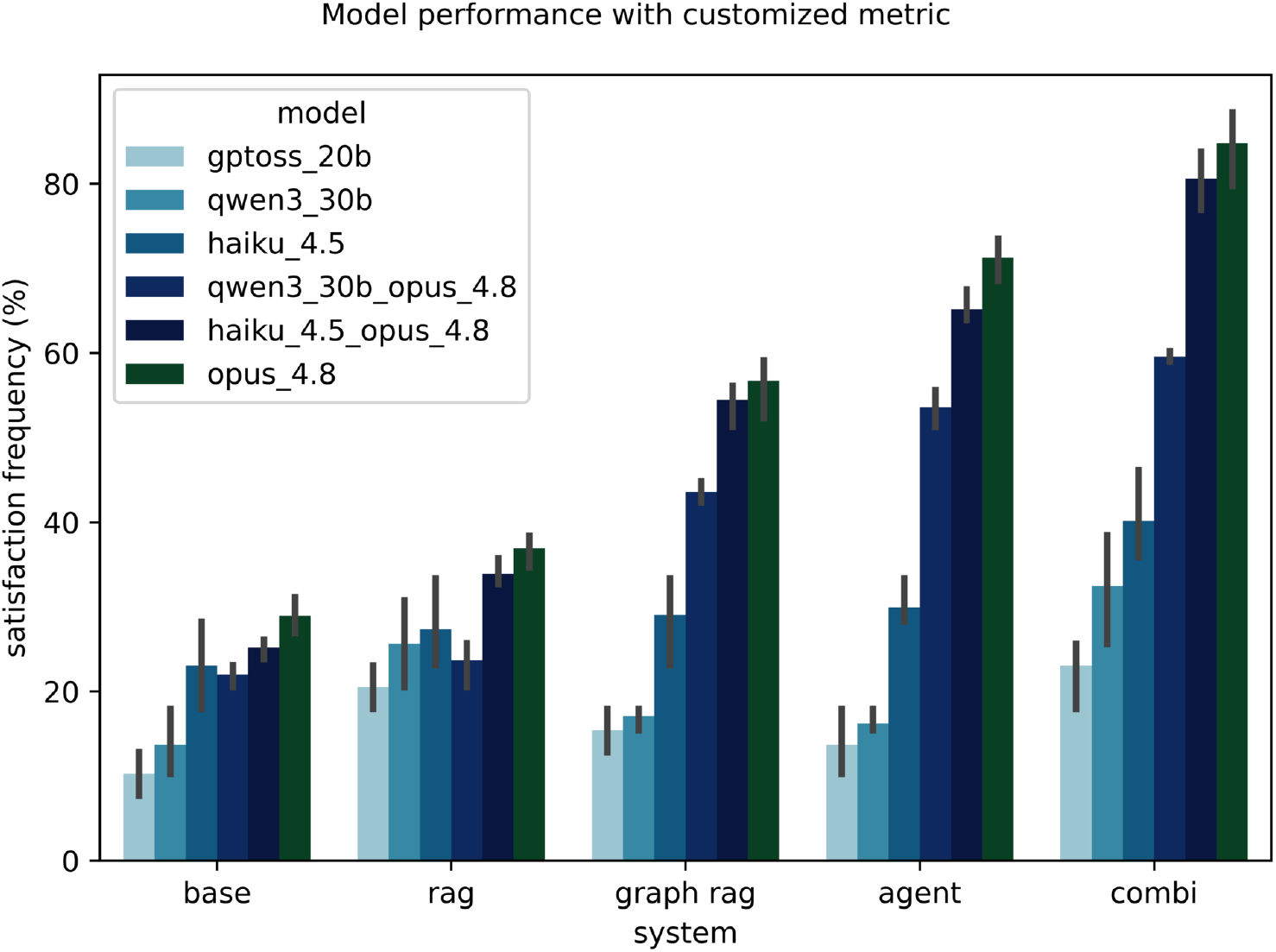
Satisfaction frequency *ψ* of the bare language model (*base*) and the four GN LLM systems, document RAG (*rag*), graph RAG (*graph*), agentic graph RAG (*agent*) and combi RAG (*combi*) on the 40-question benchmark, each run with six model configurations: the local models GPT-OSS 20B and Qwen3 30B A3B, the frontier models Claude Haiku 4.5 and Claude Opus 4.8, and two local-frontier pairs (Qwen3 30B A3B with Opus 4.8, Haiku 4.5 with Opus 4.8) that default to the cheaper model and call Opus only for hard steps. Satisfaction frequency *ψ* is the percentage of benchmark questions answered satisfactorily. Bars are medians over the 40 questions and three repeats and error bars show the spread. On the document RAG the six configurations score similarly, but on the graph, agentic and combi RAG they fan out: the local-only models stay low while Claude Opus 4.8, alone or paired, reaches the top. Both pairs recover much of the local-only shortfall, roughly doubling to tripling the satisfaction frequency *ψ* of Qwen3 30B A3B on the cognitive architectures, with the Haiku 4.5 and Opus pair ahead of the Qwen3 and Opus pair, and Opus alone still highest.

Figure 5 also indicates that Anthropic models are more useful in complex architectures (graph, agentic, combi RAG) than simple architectures (document RAG). As opposed to the base models, performance discrepancies on the document RAG with Claude Haiku 4.5, Qwen3 30B A3B and GPT-OSS 20B were minimal. In fact, Claude Haiku 4.5 and Qwen3 30B A3B had very similar satisfaction frequency *ψ* on the document RAG. The satisfaction frequency *ψ* did not vary much with Claude Haiku 4.5 but increased with Qwen3 30B A3B and GPT-OSS 20B. This suggests the effectiveness of the RAG architecture in closing the knowledge gap between the two categories of models.

In contrast, performance discrepancies on the graph RAG, agent and combi systems between Anthropic and local models were very pronounced. Systems using Claude Haiku 4.5 had twice the satisfaction frequency *ψ* of the systems leveraging Qwen3 30B A3B or GPT-OSS 20B. The gap became overwhelming with Claude Opus 4.8 (larger Anthropic model). We noted that even Claude Haiku 4.5 struggled with complex search architectures based on our RDF knowledge base. These results revealed that composing LLM with RDF requires significant cognitive capabilities. Top achievers for the graph and agentic RAG all harnessed Claude Opus 4.8.

Based on the observation that Claude Opus 4.8 drastically improved inference, we designed model combinations that could demonstrate state-of-art performance and save cost for our systems. We tried Claude Haiku 4.5 with Claude Opus 4.8 on one hand and the best local model (Qwen3 30B A3B) with Claude Opus 4.8 on the other. Typically, the system leveraging the combination of models uses the smallest model for inference by default and only exploits the capabilities of the large model for complex tasks in the pipeline. A system that uses a combination of models is called hybrid.

Figure 5 illustrates performance results of systems using those model combinations alongside single model systems. As expected, LLM systems using the combination of Qwen3 30B A3B and Claude Opus 4.8 largely surpassed LLM systems powered with Qwen3 30B A3B only, in performance. The satisfaction frequency *ψ* spiked from 17% to 43% on the graph RAG and from 16% to 53% on the agentic RAG. However, the combination of Qwen3 30B A3B and Claude Opus 4.8 exhibited a lower performance than the combination of Claude Haiku 4.5 and Claude Opus 4.8, which in turn showed a lower performance than Claude Opus 4.8 exclusively. In other words, Claude Haiku 4.5 made better use of Claude Opus 4.8 capabilities than Qwen3 30B A3B in most systems. On the combi system this ordering meant a satisfaction frequency *ψ* of about 86% with Claude Opus 4.8, 81% with the Claude Haiku 4.5 and Opus pair, and 59% with the Qwen3 30B A3B and Opus pair (Figure 5). Table 1 details expenditure for each model combination. Dividing *ψ* by cost per query shows that cost-effectiveness does not track *ψ* itself: the graph RAG returns the most *ψ* per dollar of any system (roughly 228-275 *ψ* points (%) per $1, versus 168-208 for the document RAG), while the agent and combi systems, despite scoring highest on *ψ*, return far less per dollar (37-63 and 37-51 *ψ* points (%) per $1, respectively). The combination of Qwen3 30B A3B and Claude Opus 4.8 is therefore best understood as the cheapest way to reach a given *ψ* within the agent or combi architectures specifically, not as cost-effective in absolute terms compared to the graph RAG.

**Table 1.**
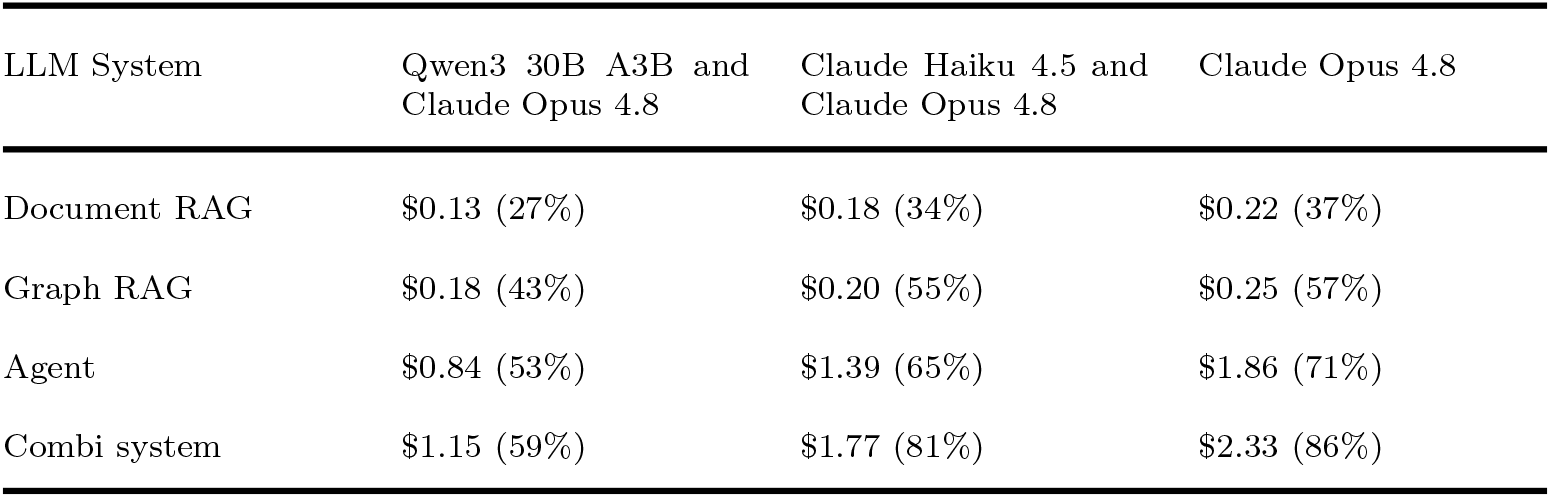
Estimated API cost per query by model combination in dollars. We recall the satisfaction frequency *ψ* for each system and combination of models in parenthesis.

## 3 Discussion

The primary objective of this study was to build an expert-curated benchmark for evaluating frontier and local LLMs querying a biomedical knowledge graph, and to use it to compare different context engineering strategies for LLM inference on scientific biomedical research. We experimented with two methods: changing the system architecture and using language models with different knowledge or training. Both techniques affect how LLMs gather context to address questions. Our findings suggest that switching to a larger or better-trained model does not always improve performance. For instance, the base Claude Opus 4.8 did not produce significantly better results than the base Claude Haiku 4.5. Similarly, Claude Haiku 4.5 did not outperform Qwen3 30B A3B on the document RAG. In fact, most LLMs used in this study, whether frontier or local, failed to generate satisfactory answers to most genetic and biomedical questions in our benchmark *on their own*. Duong and Solomon (2024) and Shang et al. (2023) emphasized similar shortcomings of LLMs on genetic questions.

System architecture, by contrast, had a substantial impact on benchmark performance. Every architecture beat the bare language model, and the agentic and graph RAG scored higher than the document RAG; with Claude Opus 4.8 the document RAG reached only 36% satisfaction frequency *ψ* against 71% for the agentic system. Similar results were reported by More (2025) and Subramanian et al. (2025). The gain was not uniform across models. We did not observe a performance boost for every model on the standalone graph or agentic systems, and only instances combined with a large model (Claude Opus 4.8) answered most questions correctly. Such behavior was also reported for other systems. For example, Nguyen et al. (2025) described a loss of benchmark performance across different agents when switching from a large to a small model. We conclude that system architecture and model together have a higher impact on performance than either alone.

The model gap was widest exactly where the architecture mattered most. On the document RAG the Anthropic and local models scored similarly, but on the graph, agentic and combi systems the local models fell well behind. Claude Haiku 4.5 beat the similarly sized local models even without any of our systems, and Qwen3 30B A3B was the stronger local model throughout. Pairing a local model with Claude Opus 4.8 recovered much of the gap, roughly doubling the satisfaction frequency *ψ* of Qwen3 30B A3B on the graph and agentic RAG, though a Claude Haiku 4.5 and Opus pair, and Opus alone, still scored higher. These two findings motivated a new system that combines the three architectures and switches models as needed. This hybrid system runs the document, graph and agentic RAG together and merges their answers. It also chooses between a small local model and a large frontier model on the fly, calling the frontier model only for the steps that require it. This gave better performance than any single architecture and lower cost than running the frontier model throughout.

Our findings are in apparent tension with Zhavoronkov et al. (2026), who reported that compact, fine-tuned Longevity-LLMs (0.6B–9B parameters) matched or exceeded far larger frontier systems on structured-omics tasks. We think this reflects a difference in setup. Their small models were fine-tuned on the target task, while our local models (Qwen3 30B A3B, GPT-OSS 20B) were generic. Also, their benchmark was tailored to the classification and regression over a fixed numeric feature set, tasks a fine-tuned small model can specialize into, whereas our graph and agentic systems require generating and revising SPARQL on the fly, on which small models struggled. It does however suggest fine-tuning a local model on SPARQL generation for GN’s schema, rather than on the target domain alone, as a promising direction.

Data representation mattered as well: the systems that queried the RDF graph directly, the graph and agentic RAG, produced our best benchmark results. This is consistent with the view that modeling data in RDF improves representation and supports reasoning over complex structures (Tomaszuk and Hyland-Wood 2020; Alemu et al. 2012; Allemang and Hendler 2011), including genetic data (Kushida et al. 2025; Sima et al. 2019; Deus et al. 2010). We built the interface with a Domain-Specific Language (DSL) that translates the GN SQL database schema to RDF, a design motivated by the closeness of RDF syntax to natural language (Tomaszuk and Hyland-Wood 2020; Gerber and Ngomo 2014) and by a growing body of work that uses RDF to improve inference with language models (Georgiadis et al. 2026; Araya et al. 2025; Perevalov and Both 2024; Regino et al. 2026). Success nonetheless depended heavily on the model. Not all LLMs could generate valid SPARQL queries, even when given a good description of the graph schema. Every small model in this work struggled with SPARQL generation, including Claude Haiku 4.5, echoing difficulties reported for small language models (Rangel et al. 2024; Frey et al. 2023). This underlines the need to improve the interface between LLMs and RDF through fine-tuning.

A further contribution of this work is the evaluation metric itself. We condensed the standard classification metrics (accuracy, precision, recall, f1) into a single score, the satisfaction frequency *ψ*, to make interpretation and comparison easier. It represents the proportion of questions for which the generated answers were similar enough to the true answers.

We encoded similarity between LLM generated and true answers as a function of the classification metrics. The function is flexible and generalizable. Depending on the use case, it could prioritize precision over recall. The function could also make use of more recent classification metrics such as cPrecision, cRecall (Yacouby and Axman 2020), informedness or markedness (Powers 2020). In this study, we noticed that some LLM generated answers were considered non-satisfactory because they contained facts not available in the true answers, affecting the precision score. To prevent such bias, we coded the function logic so that answers with good recall and fair precision or vice versa are judged satisfactory. We checked the metric against precision, recall and f1 on the local-model runs and found that both approaches ranked the systems, and the two local models, in the same way.

One methodological continuity with our earlier GNQA system (Darnell et al. 2024) is worth noting: GNQA relied on RAGAS combined with human thumbs-up/down ratings, a combination it acknowledged not always in agreement (e.g. 89% RAGAS answer relevance versus 76% human approval). We took this a step further by replacing free-form questions and RAGAS’s four-metric output with a curated, expert-authored benchmark of true answers and a single composite satisfaction frequency *ψ* derived from classification metrics, aimed at making the standard versus AI evaluation gap identified in GNQA easier to close and to interpret. We also replaced the closed FahamuAI’s stack with DSPy and LangChain, both open-source frameworks for LLM system design.

We acknowledge the statistical limitations of this work in terms of sample representation. Evaluating performance over more iterations followed by statistical testing would give a clearer picture of how context engineering strategies affect LLM inference and cost. Looking at the results we decided that the runs we did were representative enough for our purposes. In addition to more runs the analysis can be extended to a wider selection of both frontier and local models (there are hundreds), and the benchmark expanded to cover more questions in medical genetics.

Our results point to a useful division of labour for the hybrid systems: a small local model for routine steps and orchestration, and a larger frontier model for the parts that need sophisticated parsing and reasoning, such as SPARQL generation and answer synthesis. This keeps most inference local and inexpensive while reserving the frontier model for where it is needed.

Finally, the AI industry is moving fast and anything we report will be out-of-date by the time it gets written up. Even so, we attempted to come up with a method and report that should be relevant for the coming years. There are good reasons for using local models, such as cost and privacy. There are issues with reproducibility between individual frontier model updates. Worse, in recent developments, frontier models are blocking queries related to biology. In contrast, smaller models can be downloaded and queried locally and we show results are fairly reproducible. In future work, we may train our own specialized small models for tasks such as SPARQL query construction over the data structures we have in GN, shifting more of that load off the frontier model while allowing improved reproducibility and control. We intend to develop our hybrid/combi solution further, teaching it to select subsystems by task rather than running all three in parallel, which would cut computational cost while preserving answer quality, and to widen the set of models it can route between according to task complexity. We are also interested in speeding up the systems. This can potentially be achieved by improving the hardware, by caching, and by running faster models. For exploring users it will be very interesting to get the full response in a scale of seconds rather than minutes.

## 4 Methods

### 4.1 We rebuilt the GN metadata layer as an RDF graph

GN stored its data in a MariaDB relational database that became hard to navigate and slow to query as it grew in size and complexity (Babucea 2021; Kleppmann 2017; Chen and Zhang 2014). Graph technologies capture complex biological relationships well (Sima et al. 2019; Deus et al. 2010) and key-value stores retrieve large datasets quickly (Agrawal and Mamajiwala 2024; Lin et al. 2024; Slabinoha et al. 2022). We therefore re-modeled the data as an RDF graph queried through SPARQL(Gaignard et al. 2023; Poveda-Villalón et al. 2020), and kept the bulk phenotype matrices in an LMDB key-value store for fast access (Agrawal and Mamajiwala 2024).

#### 4.1.1 GN data lived in a 1 TB MariaDB schema of 93 interlinked tables

At the time of writing the GN database held about 1 TB, reflecting a research focus that has broadened over the years from central nervous system mouse genetics to many organisms and topics (Ashbrook et al. 2021; Watson and Ashbrook 2020; Mulligan et al. 2017; Lu and Lu 2013; Druka et al. 2008; Williams 1994). Its 93 interlinked tables cross-reference genotype and marker data, classical phenotypes and expression phenotypes (microarray or mRNA), and the schema is hard to use without prior knowledge of it (Watson and Ashbrook 2020; Sloan et al. 2016; Mulligan et al. 2017).

#### 4.1.2 A DSL maps the SQL schema to RDF served over a SPARQL REST API

We wrote a self-documenting Domain-Specific Language (DSL) in GNU Guile (v.5.0.1) that maps SQL relationships to RDF triples (Figure 6). A *define-transformer* form combines a SQL query, a SQL-to-RDF predicate mapping and hand-written triples for concepts the SQL design does not cover; a *transform-with-documentation* form runs one or more transformers (not to be confused with LLM transformers) with a set of RDF prefixes and emits a Turtle file plus its Markdown documentation (Appendices A and B). We validated the Turtle with rapper (raptor v.2.0.16), loaded it into Virtuoso (v.7.2.15), and refined the schema iteratively with GN’s biologists. The graph is browsable through a resource classification scheme^1^ and a public SPARQL endpoint^2^.

**Fig. 6.**
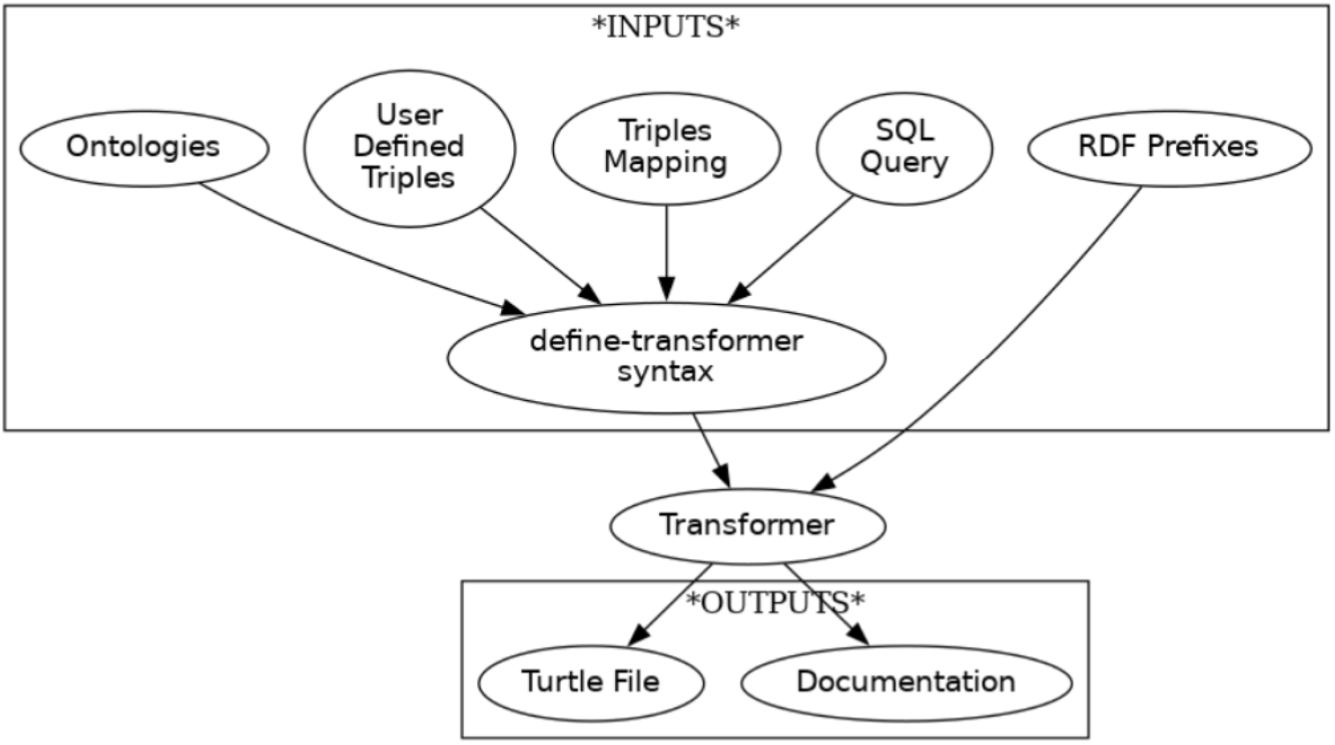
How the Domain-Specific Language turns the GN SQL schema into RDF. A *define-transformer* form ties together the SQL query, the SQL-to-RDF triple mapping, user-defined triples and the ontologies it draws on. The *transform-with-documentation* form then takes one or more such transformers plus the RDF prefixes, runs the transformation and emits a Turtle file and its Mark-down documentation.

Individual phenotype datasets are too large for RDF, so we exported them from MariaDB to LMDB (v.0.9.29)^3^, keyed by trait, storing every individual or strain measurement. An API endpoint^4^ returns a trait’s measurements as JSON, and its parameterised URL is stored in the RDF graph so experimental data can be reached directly from a SPARQL result. All software was pinned with GNU Guix (v.1.5.0).

### 4.2 We built four LLM systems over the RDF graph and literature corpus

LLMs are increasingly used to navigate and reason over biomedical data (Huang et al. 2025; Jin et al. 2025; Luo et al. 2025; Aamer et al. 2025; Zhou et al. 2025; Lan et al. 2025; Ifargan et al. 2025; Bi et al. 2024; Lu et al. 2024; Wysocki et al. 2024). GN’s many datasets have been hard to discover and reuse; the RDF graph makes them queryable, and on top of it we built four LLM systems that answer questions about GN data through a chat interface. The systems are a document RAG, a graph RAG, an agentic graph RAG and a combination of the three called ‘combi’; Table 2 compares them and sections 4.2.1–4.2.4 describe each.

**Table 2.**
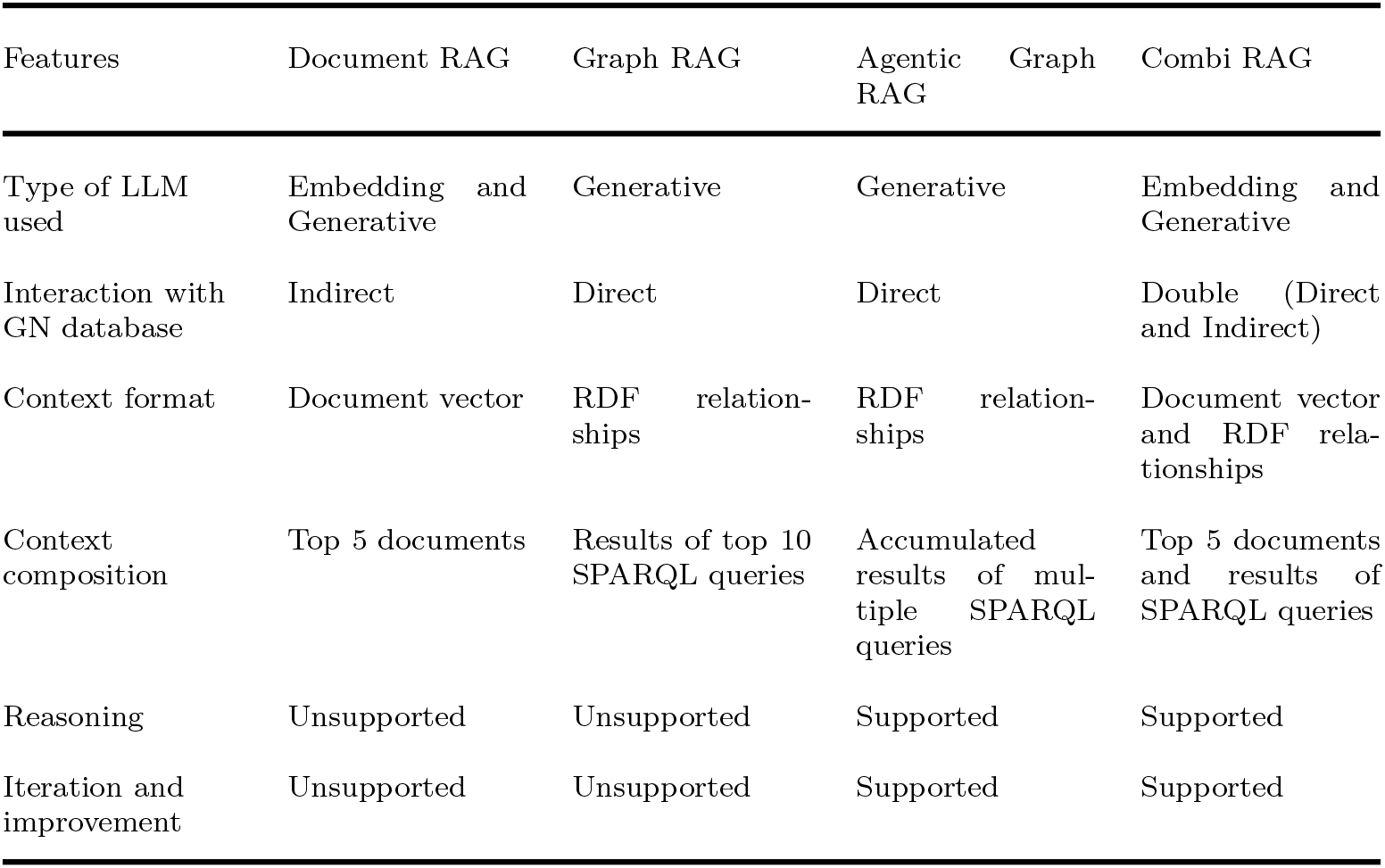
Comparison of GN LLM Systems.

#### 4.2.1 The document RAG answers from text passages retrieved by keyword and embedding search

This is the standard RAG design: retrieve text passages relevant to the question and let the LLM answer from them (Fan et al. 2024; Arslan et al. 2024; Gupta et al. 2024; Abo El-Enen et al. 2025; Singh et al. 2025). We built the corpus from the RDF graph: targeted SPARQL queries^5^ pulled model organism classifications, phenotypes, genotypes and gene annotations (NCBI GeneRIF, NCBI wiki), a hand-curated predicate-to-verb mapping turned the triples into plain-text sentences, and sentences sharing a subject were grouped into one document per subject (Figure 7). At query time the question is embedded with a Qwen3 embedding model^6^ and matched against a Chroma vector store by cosine similarity, and in parallel a BM25 keyword search is run; the two result sets are combined with query-dependent weights. The corpus is indexed and retrieved with LangChain (v.1.2.10) (Chase 2022), the LLM interface is managed with DSPy (v.3.1.3) (Khattab et al. 2023), and the code is on GitHub^7^.

**Fig. 7.**
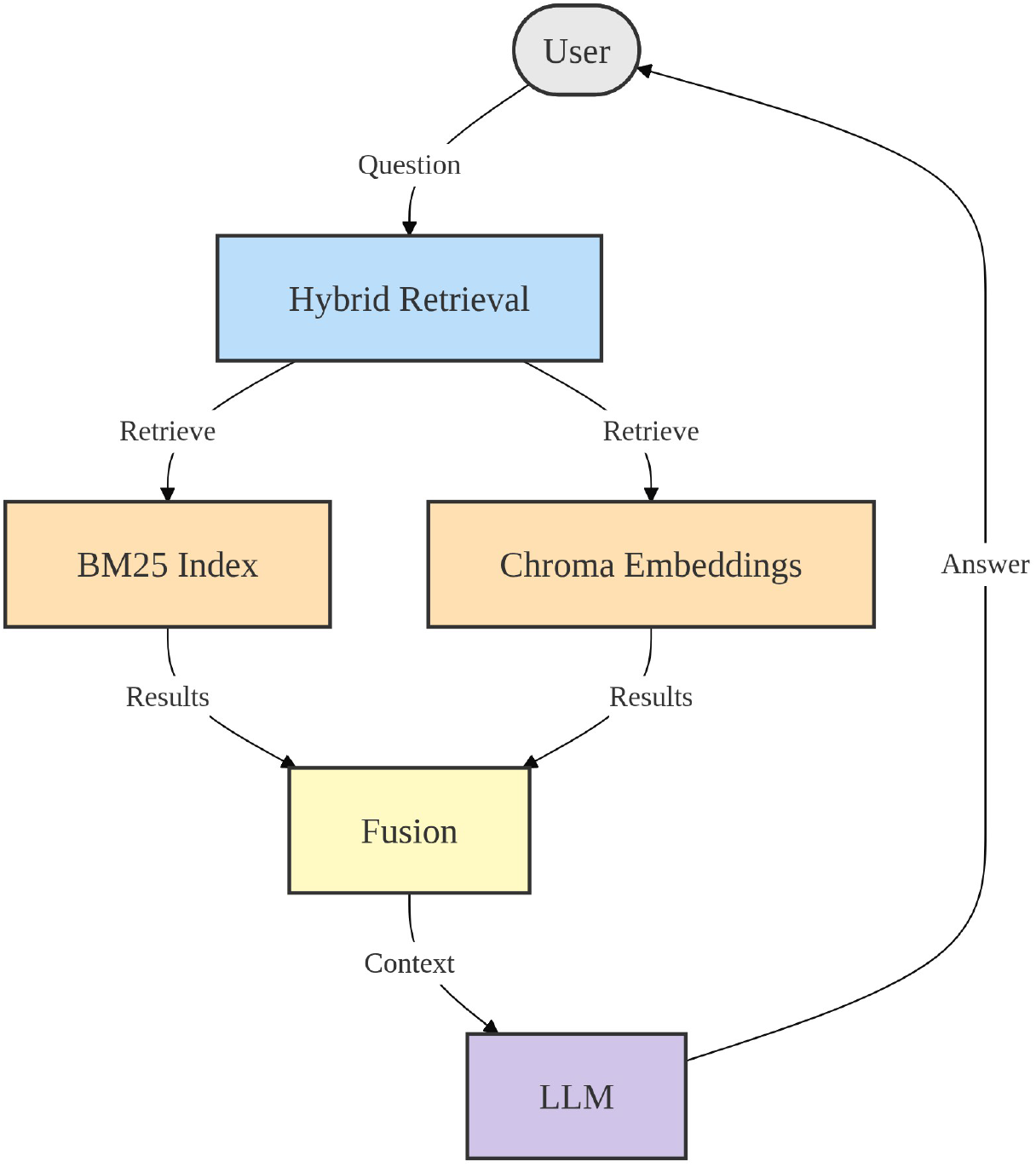
How the document RAG answers a question. A text corpus is curated from the RDF database and indexed both with BM25 (keyword) and as embeddings in a Chroma database. The user question is run against both indexes, the two result sets are fused into a single context, and the LLM generates the answer from that context.

#### 4.2.2 The graph RAG answers from the results of LLM-generated SPARQL queries

Instead of pre-converting the graph to text, the graph RAG lets the LLM query it directly (Smeros et al. 2025; Rangel et al. 2024; Chen et al. 2021; Singh et al. 2025; Zou et al. 2014) (Figure 8). It has two modules: an RDF-aware LLM that generates ten SPARQL queries for the endpoint from the question and schema hints (RDF classes, properties and example triples), and a second LLM that writes the answer from the returned rows. Keywords extracted from the question are matched against the hints to guide query generation. Both modules are built with DSPy (v.3.1.3); we did not use federation across endpoints.

**Fig. 8.**
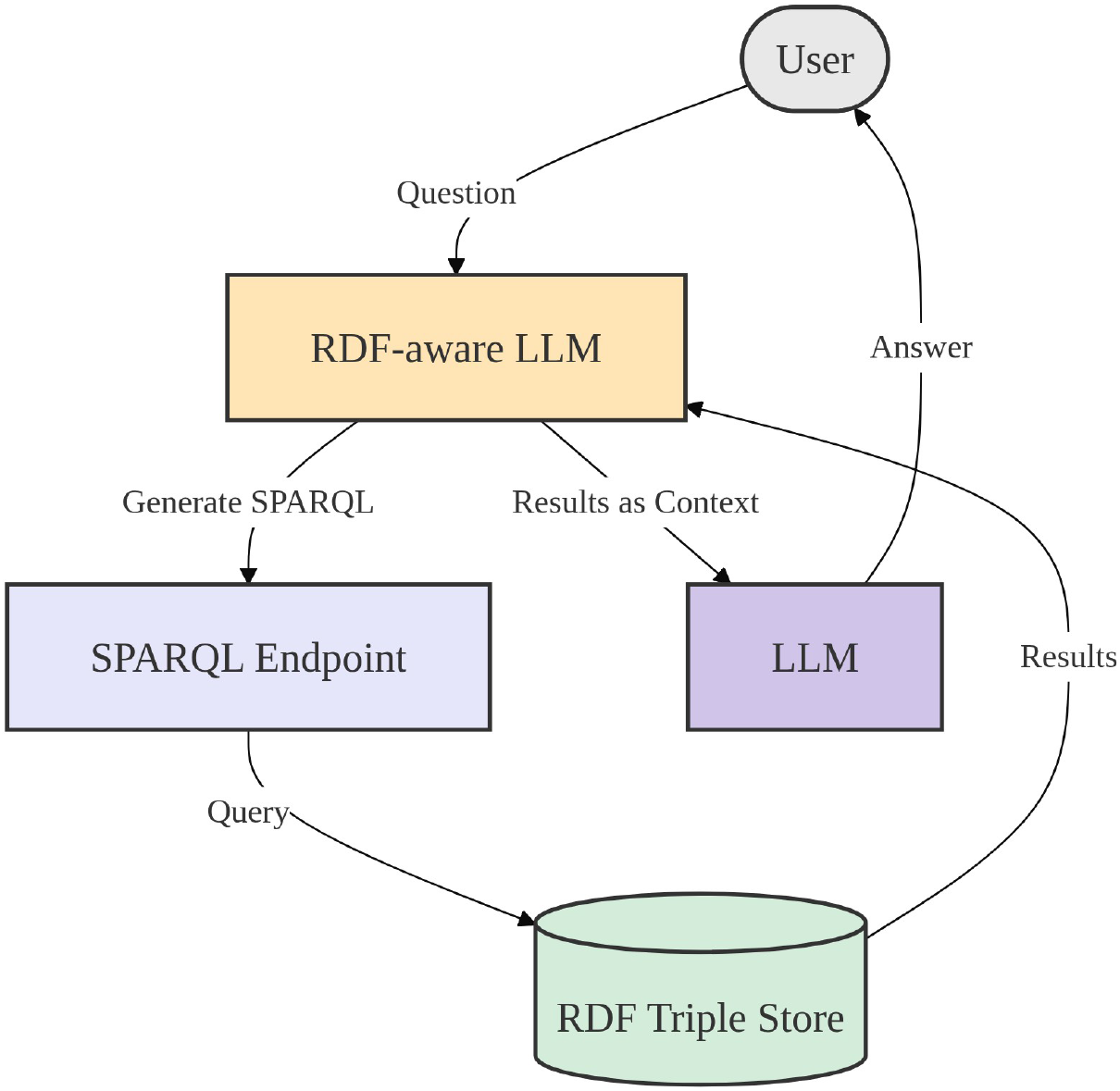
How the graph RAG answers a question. An RDF-aware LLM turns the question into a set of SPARQL queries, using the classes, properties and example triples of the graph schema as hints. The queries are run against the SPARQL endpoint over the RDF triple store, and the returned rows become the context a second LLM call uses to write the answer.

#### 4.2.3 The agentic graph RAG refines its SPARQL over up to 20 ReAct cycles

The graph RAG queries the endpoint once, which often misses context. The agentic graph RAG instead runs a ReAct loop (Singh et al. 2025): the agent generates a SPARQL query, inspects the results and revises or extends it, for up to 20 reason-act-observe cycles, before writing the answer from everything it has gathered (Figure 9). It is implemented with DSPy (v.3.1.3); the code is on GitHub^8^.

**Fig. 9.**
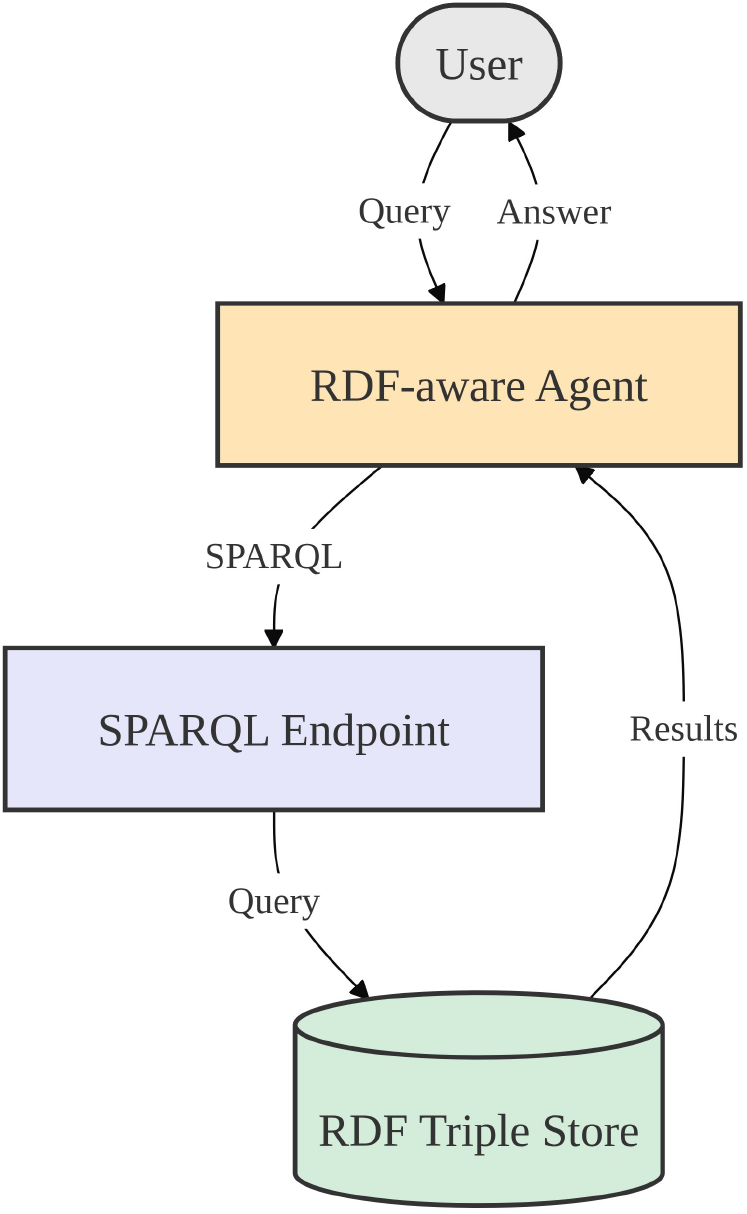
How the agentic graph RAG answers a question. An RDF-aware agent sends a SPARQL query to the endpoint, inspects the returned rows and revises or extends the query. It repeats this reason-act-observe loop for up to 20 ReAct cycles, accumulating context, then writes the answer from everything it has gathered.

#### 4.2.4 The Combi RAG runs all three systems concurrently and merges their answers

The combi RAG runs the document, graph and agentic graph RAG in parallel, streams their intermediate outputs, and synthesises a single answer from the three (Figure 10). Conversation memory is kept between turns with mem0ai (v.2.0.1) (Chhikara et al. 2025), the system is built with DSPy (v.3.1.3) (Khattab et al. 2023), and the code is on GitHub.

**Fig. 10.**
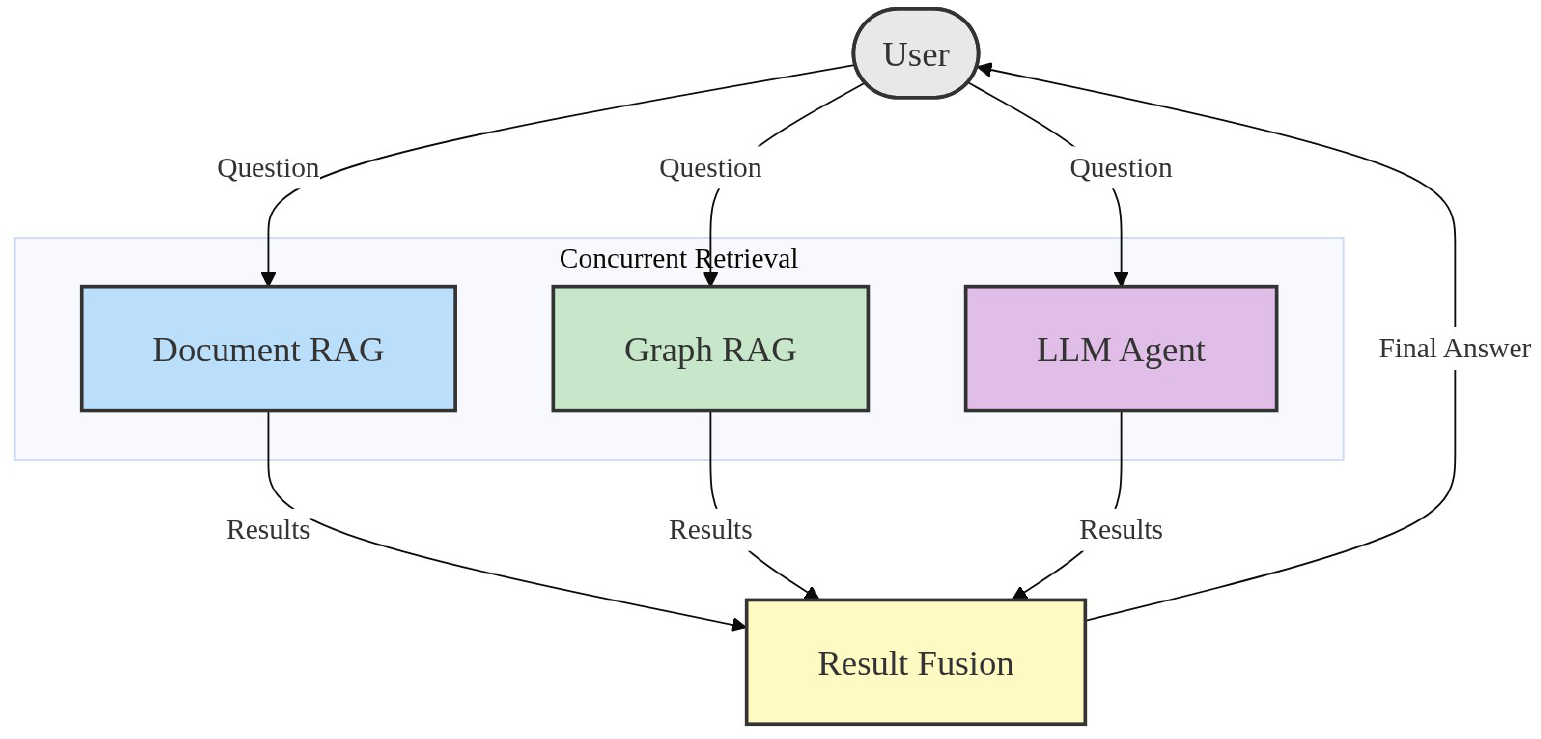
How the combi RAG answers a question. The question is sent to the document RAG, graph RAG and agentic graph RAG at the same time, and each streams its own output. A fusion step then synthesises the three results into the final answer, and conversation memory is kept between queries so follow-ups can build on earlier turns.

### 4.3 We evaluate four architectures across six model configurations

We evaluated the four architectures on a fixed benchmark while varying the language model behind them: four single models and two local-frontier pairs (Table 3). Sections 4.3.2 and 4.3.3 give the metric and the scoring procedure.

**Table 3.** Performance Evaluation Design.

| Model selection | Run ID | Document RAG | Graph RAG | Agentic Graph RAG | Combi RAG |
| --- | --- | --- | --- | --- | --- |
| Single | Run 1 | Claude 4.8 Opus | Claude 4.8 Opus | Claude 4.8 Opus | Claude 4.8 Opus |
|  | Run 2 | Claude 4.5 Haiku | Claude 4.5 Haiku | Claude 4.5 Haiku | Claude 4.5 Haiku |
|  | Run 3 | Qwen 3 A3B | Qwen 3 A3B | Qwen 3 A3B | Qwen 3 A3B |
|  | Run 4 | GPT OSS | GPT OSS | GPT OSS | GPT OSS |
| Pair | Run 5 | Claude 4.5 Haiku and Claude 4.8 Opus | Claude 4.5 Haiku and Claude 4.8 Opus | Claude 4.5 Haiku and Claude 4.8 Opus | Claude 4.5 Haiku and Claude 4.8 Opus |
|  | Run 6 | Qwen 3 A3B and Claude 4.8 Opus | Qwen 3 A3B and Claude 4.8 Opus | Qwen 3 A3B and Claude 4.8 Opus | Qwen 3 A3B and Claude 4.8 Opus |

#### 4.3.1 Experts curated 40 questions and reference answers from the resource classification scheme

GN’s resource classification scheme provided ground-truth labels from which domain experts curated 40 questions and reference answers of varying difficulty, covering model organisms, datasets and mapping results (Mohammadi et al. 2025). The benchmark is on GitHub^9^. This benchmark was drawn from the same resource classification scheme used while developing the RDF graph and the four systems, and there is no independent held-out test set; we cannot rule out that iterating on our own classification scheme introduced some bias.

#### 4.3.2 We score answers with the satisfaction frequency *ψ*, built from precision, recall and f1

Prior work scores biomedical LLM answers with precision, recall and f1 computed from an item-level confusion matrix (Qu et al. 2026; Huang et al. 2025; Obi 2023; Riyanto et al. 2023; Naidu et al. 2023). We found these four numbers hard to read together, and sometimes unfair: an LLM judge would mark a correct answer down for stating relevant facts absent from the reference, lowering precision, regardless of which model judged. We therefore defined a single *satisfaction frequency ψ*: a response counts as satisfactory when precision and recall are both at least 70 %, or f1 is at least 60 %, and the satisfaction frequency *ψ* is the fraction of benchmark questions a system answers satisfactorily. Despite its name, *ψ* is a fixed-threshold summary of the LLM judge’s precision/recall/f1 scores, not a measure calibrated against actual user satisfaction judgments; we did not run a human study to validate the 70%/60% thresholds or the judge’s scoring against human raters, so *ψ* should be read as a proxy for the standard classification metrics rather than a validated user satisfaction metric. For the comparison in section 2.1 we computed both the standard averages^10^ and the satisfaction frequency *ψ*^11^ on the local-model runs, with plots^12^ in matplotlib (v.3.11.0).

#### 4.3.3 An LLM judge scores each answer over three runs to compare architectures

Each system answered every benchmark question with up to five retries, using the same model across systems, and we also ran each model on its own as a baseline (Sharma and Mehta 2025; Lei et al. 2025; de Aquino e Aquino et al. 2025). A single LLM judge (Claude Haiku 4.5) compared each answer to its reference and counted true positives (statements in both), false positives (only in the answer) and false negatives (only in the reference); true negatives cannot be defined here, so accuracy is not used (Qu et al. 2026; Huang et al. 2025). We did not check this judge’s consistency against a second judge or human raters, so judge-specific bias cannot be ruled out. From these, precision is TP*/*(TP + FP), recall is TP*/*(TP + FN) and f1 is 2 TP*/*(2 TP + FP + FN), which feed the satisfaction frequency *ψ* of section 4.3.2. We repeated the whole evaluation three times at temperature 1 with caching disabled, and also recorded wall-clock time and token count per query; plots^13^ use matplotlib (v.3.11.0). Each of the three repeats scores *ψ* once over the full 40-question benchmark, so *N* = 3 is the sample size for statistical comparisons between systems or models; the per-question judgments underlying each repeat’s *ψ* are not retained separately, which limits us to paired tests across repeats rather than across questions. We report paired t-tests on these three repeats where we compare two conditions directly (sections 2.2 and 2.4); with only two degrees of freedom such tests have limited power, so a non-significant result should not be read as evidence of no difference.

### 4.4 We test frontier models, local models and local-frontier pairings on the same systems

Model capability matters as much as architecture, and open local models are far cheaper than frontier ones (Safavi-Naini et al. 2025; Mavrych et al. 2025; Pan et al. 2025). As frontier models we used Claude Opus 4.8 and Claude Haiku 4.5; as local models, Qwen3 30B A3B Thinking 2507 (Q4 K M)^14^ and GPT-OSS 20B (F16)^15^, served with ollama (v.0.31.1) on an RTX 3090 (24 Gb VRAM). Haiku 4.5 is the size-matched comparator for the local models and Opus 4.8 is the state-of-the-art reference. We also tested two mixed configurations, Haiku 4.5 with Opus 4.8 and Qwen3 with Opus 4.8, each defaulting to the smaller model and calling Opus only for the hard steps of the pipeline; all six configurations are listed in Table 3 and evaluated as in section 4.3.3.

## Supplementary information

## Acknowledgements

We acknowledge Dr Suhaila Hashim and Pwani University for providing space and facilities. We acknowledge the Sambamba Foundation for providing computational resources. Claude (Anthropic) was used to check clarity and structure during editing; no text in this manuscript was drafted unchecked by AI.

Claude Opus 4.8 and Claude Haiku 4.5 (Anthropic) were used as the frontier models in our experiments. The local models Qwen3 30B A3B (Alibaba Cloud/Qwen team) and GPT-OSS 20B (OpenAI) are gratefully acknowledged.

## Declarations

The authors have declared no competing interest. All experiments with Qwen were performed in Kenya for demonstration purposes only.

## Appendix A Appendix

**Fig. A1.**
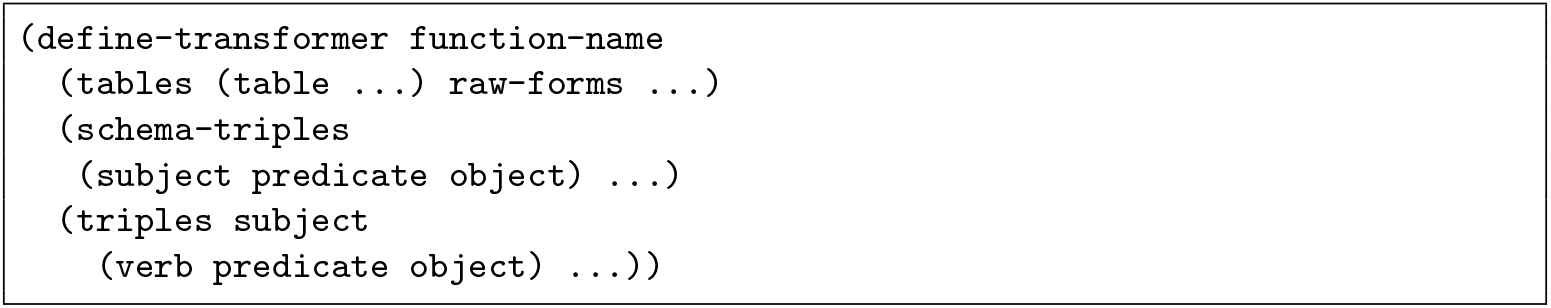
The *define-transformer* form declares one SQL-to-RDF transformation. It has three order-agnostic clauses. The *tables* clause names the database tables joined to build the SQL view. The *schema-triples* clause lists triples written once for the whole view. The *triples* clause lists triples emitted once per row of the view.

## Appendix B Appendix

**Fig. B2.**
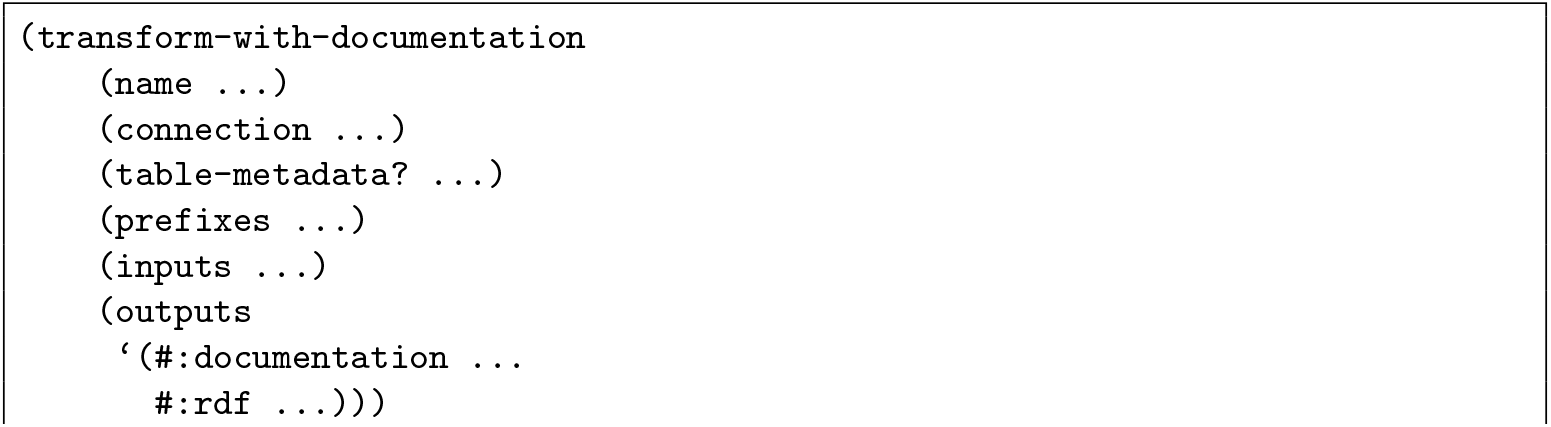
The *transform-with-documentation* form runs a set of transformers and writes the results. *name* is the title of the generated Markdown file. *connection* holds the SQL connection settings. *table-metadata?* toggles inclusion of table-level metadata. *prefixes* are the ontology prefixes placed at the top of the Turtle file. *inputs* lists the *define-transformer* forms to execute. *outputs* gives the paths for the documentation and RDF files.

## Footnotes

1 https://rdf.genenetwork.org/v1/category/resource_classification_scheme

2 https://rdf.genenetwork.org/sparql

3 https://github.com/genenetwork/genenetwork3/blob/main/scripts/lmdb_phenotypes_matrix.py

4 https://github.com/genenetwork/genenetwork3/blob/main/gn3/api/lmdb_traits.py

5 https://git.genenetwork.org/genenetwork_ai_search/tree/aisearch/scripts/fetch_metadata.py

6 https://huggingface.co/Qwen/Qwen3-Embedding-0.6B

7 https://git.genenetwork.org/genenetwork_ai_search/tree/aisearch

8 https://git.genenetwork.org/genenetwork_ai_search/tree/aisearch/src/gnais/search/agent.py

9 https://git.genenetwork.org/genenetwork_ai_search/tree/aisearch/data/small_benchmark.csv

10 https://git.genenetwork.org/genenetwork_ai_search/tree/aisearch/scripts/simple_evaluate.py

11 https://git.genenetwork.org/genenetwork_ai_search/tree/aisearch/scripts/dspy_evaluate.py

12 https://git.genenetwork.org/genenetwork_ai_search/tree/aisearch/scripts/barplot_metrics.py

13 https://git.genenetwork.org/genenetwork_ai_search/tree/aisearch/scripts/plot_function.py

14 https://huggingface.co/unsloth/Qwen3-30B-A3B-Thinking-2507-GGUF

15 https://huggingface.co/unsloth/gpt-oss-20b-GGUF/blob/main/gpt-oss-20b-F16.gguf

## References

Aamer N, Asim MN, Munir S, et al (2025) Automating AI discovery for biomedicine through knowledge graphs and LLM agents. bioRxiv pp 2025–05. 10.1101/2025.05.08.652829

Abo El-Enen M, Saad S, Nazmy T (2025) A survey on retrieval-augmentation generation (RAG) models for healthcare applications. Neural Computing and Applications 37(33):28191–28267. 10.1007/s00521-025-11666-9

Agrawal A, Mamajiwala Y (2024) Comparative analysis of rocksdb, lmdb, and mon-godb: A performance evaluation. Tech. rep., SAE Technical Paper, 10.4271/2024-26-0331

Alemu G, Stevens B, Ross P, et al (2012) Linked data for libraries: benefits of a conceptual shift from library-specific record structures to RDF-based data models. New library world 113(11-12):549–570. 10.1108/03074801211282920

Allemang D, Hendler J (2011) Semantic web for the working ontologist: effective modeling in RDFS and OWL. Elsevier

de Aquino e Aquino G, de Azevedo NdS, Okimoto LYS, et al (2025) From rag to multi-agent systems: A survey of modern approaches in llm development. Preprints, 10.20944/preprints202502.0406.v1

Araya D, Valle C, Astudillo H, et al (2025) Enhancing contextual memory in LLMs for software engineering via ontology-based inference. In: WAAI/AIESD/WDEA/WKMIT/SCTSD/WSM, pp 45–54

Arslan M, Ghanem H, Munawar S, et al (2024) A survey on RAG with LLMs. Procedia computer science 246:3781–3790. 10.1016/j.procs.2024.09.178

Ashbrook DG, Arends D, Prins P, et al (2021) A platform for experimental precision medicine: The extended bxd mouse family. Cell systems 12(3):235–247. 10.1016/j.cels.2020.12.002

Babucea AG (2021) Sql or NoSQL databases? critical differences. Annals of ‘Constantin Brancusi’ University of Targu-Jiu

Bi Z, Dip SA, Hajialigol D, et al (2024) Ai for biomedicine in the era of large language models. arXiv preprint arXiv:240315673 10.48550/arXiv.2403.15673

Chase H (2022) LangChain. URL https://github.com/langchain-ai/langchain

Chen CP, Zhang CY (2014) Data-intensive applications, challenges, techniques and technologies: A survey on big data. Information sciences 275:314–347. 10.1016/j.ins.2014.01.015

Chen YH, Lu EJL, Ou TA (2021) Intelligent SPARQL query generation for natural language processing systems. IEEE Access 9:158638–158650. 10.1109/ACCESS.2021.3130667

Chhikara P, Khant D, Aryan S, et al (2025) Mem0: Building production-ready AI agents with scalable long-term memory. arXiv preprint arXiv:250419413 10.48550/arXiv.2504.19413

Darnell SS, Overall RW, Guarracino A, et al (2024) Creating a biomedical knowledge base by addressing gpt inaccurate responses and benchmarking context. bioRxiv p 2024.10.16.618663. 10.1101/2024.10.16.618663, gNQA: GeneNetwork Question-Answer knowledge base

Deus HF, Veiga DF, Freire PR, et al (2010) Exposing the cancer genome atlas as a SPARQL endpoint. Journal of biomedical informatics 43(6):998–1008. 10.1016/j.jbi.2010.09.004

Druka A, Druka I, Centeno AG, et al (2008) Towards systems genetic analyses in barley: Integration of phenotypic, expression and genotype data into genenetwork. BMC genetics 9(1):73. 10.1186/1471-2156-9-73

Duong D, Solomon BD (2024) Analysis of large-language model versus human performance for genetics questions. European Journal of Human Genetics 32(4):466–468. 10.1038/s41431-023-01396-8

Fan W, Ding Y, Ning L, et al (2024) A survey on rag meeting llms: Towards retrieval-augmented large language models. In: Proceedings of the 30th ACM SIGKDD conference on knowledge discovery and data mining, pp 6491–6501, 10.1145/3637528.3671470

Frey J, Meyer LP, Arndt N, et al (2023) Benchmarking the abilities of large language models for RDF knowledge graph creation and comprehension: how well do LLMs speak turtle? arXiv preprint arXiv:230917122 10.48550/arXiv.2309.17122

Gaignard A, Rosnet T, De Lamotte F, et al (2023) Fair-checker: supporting digital resource findability and reuse with knowledge graphs and semantic web standards. Journal of Biomedical Semantics 14(1):7. 10.1186/s13326-023-00289-5

Georgiadis T, Pavlopoulos J, Mamoulis N (2026) Sparagraph: Spatial reasoning using retrieval-augmented generation. ACM Transactions on Spatial Algorithms and Systems 12(3):1–31. 10.1145/3799424

Gerber D, Ngomo ACN (2014) From RDF to natural language and back. In: Towards the Multilingual Semantic Web: Principles, Methods and Applications. Springer, p 193–209, 10.1007/978-3-662-43585-4_12

Gupta S, Ranjan R, Singh SN (2024) A comprehensive survey of retrieval-augmented generation (rag): Evolution, current landscape and future directions. arXiv preprint arXiv:241012837 10.48550/arXiv.2410.12837

Huang K, Zhang S, Wang H, et al (2025) Biomni: A general-purpose biomedical ai agent. biorxiv 10.1101/2025.05.30.656746

Ifargan T, Hafner L, Kern M, et al (2025) Autonomous LLM-driven research—from data to human-verifiable research papers. NEJM AI 2(1):AIoa2400555. 10.1056/AIoa2400555

Jin R, Zhang Z, Wang M, et al (2025) Stella: Self-evolving LLM agent for biomedical research. 10.48550/arXiv.2507.02004, arXiv:2507.02004

Khattab O, Singhvi A, Maheshwari P, et al (2023) Dspy: Compiling declarative language model calls into self-improving pipelines. URL https://arxiv.org/abs/2310.03714, arXiv:2310.03714

Kleppmann M (2017) Designing data-intensive applications: The big ideas behind reliable, scalable, and maintainable systems. O’Reilly Media, Inc.

Kushida T, de Farias TM, Sima AC, et al (2025) Federated SPARQL query performance evaluation for exploring disease model mouse: combining gene expression, orthology, and disease knowledge graphs. BMC medical informatics and decision making 25(Suppl 1):189. 10.1186/s12911-025-03013-8

Lan W, Tang Z, Liu M, et al (2025) The large language models on biomedical data analysis: A survey. IEEE Journal of Biomedical and Health Informatics 29(6):4486–4497. 10.1109/JBHI.2025.3530794

Lei Y, Xu J, Liang CX, et al (2025) Large language model agents: A comprehensive survey on architectures, capabilities, and applications. Preprints, 10.20944/preprints202512.2119.v1

Lin H, Wang Z, Qi S, et al (2024) Building a high-performance graph storage on top of tree-structured key-value stores. Big Data Mining and Analytics 7(1):156–170. 10.26599/BDMA.2023.9020015, URL 10.26599/bdma.2023.9020015

Lu H, Lu L (2013) Genetic dissection of pax6 through genenetwork. In: Frontier and Future Development of Information Technology in Medicine and Education: ITME 2013. Springer, p 3151–3156, 10.1007/978-94-007-7618-0_404

Lu Z, Peng Y, Cohen T, et al (2024) Large language models in biomedicine and health: current research landscape and future directions. Journal of the American Medical Informatics Association 31(9):1801–1811. 10.1093/jamia/ocae202

Luo Y, Shi L, Li Y, et al (2025) From intention to implementation: automating biomedical research via LLMs. Science China Information Sciences 68(7):170105. 10.1007/s11432-024-4485-0

Mavrych V, Yaqinuddin A, Bolgova O (2025) Claude, chatgpt, copilot, and gemini performance versus students in different topics of neuroscience. Advances in Physiology Education 49(2):430–437. 10.1152/advan.00093.2024

Mohammadi M, Li Y, Lo J, et al (2025) Evaluation and benchmarking of llm agents: A survey. In: Proceedings of the 31st ACM SIGKDD Conference on Knowledge Discovery and Data Mining V. 2, pp 6129–6139, 10.1145/3711896.3736570

More R (2025) A unified evaluation framework for grounded LLM architectures: Comparative analysis of RAG, Self-RAG, and agentic RAG. In: 2025 5th International Conference on Artificial Intelligence and Signal Processing (AISP), IEEE, pp 1–5

Mulligan MK, Mozhui K, Prins P, et al (2017) Genenetwork: a toolbox for systems genetics. In: Systems genetics: Methods and protocols. Springer, p 75–120, 10.1007/978-1-4939-6427-7_4

Naidu G, Zuva T, Sibanda EM (2023) A review of evaluation metrics in machine learning algorithms. In: Computer science on-line conference, Springer, pp 15–25, 10.1007/978-3-031-35314-7_2

Nguyen T, Chin P, Tai YW (2025) Ma-rag: Multi-agent retrieval-augmented generation via collaborative chain-of-thought reasoning. arXiv preprint arXiv:250520096 10.48550/arXiv.2505.20096

Obi JC (2023) A comparative study of several classification metrics and their performances on data. World Journal of Advanced Engineering Technology and Sciences 8(1):308–314. 10.30574/wjaets.2023.8.1.0054

Pan G, Chodnekar V, Roy A, et al (2025) A cost-benefit analysis of on-premise large language model deployment: Breaking even with commercial LLM services. arXiv preprint arXiv:250918101 10.48550/arXiv.2509.18101

Parker CC, Dickson PE, Philip VM, et al (2017) Systems genetic analysis in genenetwork. org. Current protocols in neuroscience 79(1):8–39. 10.1002/cpns.23

Perevalov A, Both A (2024) Towards LLM-driven natural language generation based on SPARQL queries and RDF knowledge graphs. In: Proceedings of the Joint Proceedings of the 3rd International Workshop on Knowledge Graph Generation from Text (TEXT2KG) and Data Quality Meets Machine Learning and Knowledge Graphs (DQMLKG) Co-Located with the Extended Semantic Web Conference (ESWC 2024), Hersonissos, Greece, pp 26–30

Poveda-Villalón M, Espinoza-Arias P, Garijo D, et al (2020) Coming to terms with FAIR ontologies. In: International Conference on Knowledge Engineering and Knowledge Management, Springer, pp 255–270, 10.1007/978-3-030-61244-3_18

Powers DMW (2020) Evaluation: from precision, recall and f-measure to ROC, informedness, markedness and correlation. 10.48550/arXiv.2010.16061, URL https://arxiv.org/abs/2010.16061, arXiv:2010.16061

Qu Y, Huang K, Yin M, et al (2026) Crispr-GPT for agentic automation of geneediting experiments. Nature Biomedical Engineering 10(2):245–258. 10.1038/s41551-025-01463-z

Rangel JC, de Farias TM, Sima AC, et al (2024) Sparql generation: an analysis on fine-tuning openllama for question answering over a life science knowledge graph. arXiv preprint arXiv:240204627 10.48550/arXiv.2402.04627

Regino AG, Rossanez A, Torres RdS, et al (2026) A systematic literature review on RDF triple generation from natural language texts. Semantic Web 17(1):22104968251398355. 10.1177/22104968251398355

Riyanto S, Imas SS, Djatna T, et al (2023) Comparative analysis using various performance metrics in imbalanced data for multi-class text classification. International Journal of Advanced Computer Science and Applications 14(6). 10.14569/IJACSA.2023.01406116

Safavi-Naini SAA, Ali S, Shahab O, et al (2025) Benchmarking proprietary and open-source language and vision-language models for gastroenterology clinical reasoning. npj Digital Medicine 10.1038/s41746-025-02174-0

Shang X, Liao X, Ji Z, et al (2023) Benchmarking large language models for genomic knowledge with GeneTuring. BioRxiv: The Preprint Server for Biology 10.1101/2023.03.11.532238

Sharma R, Mehta M (2025) Small language models for agentic systems: A survey of architectures, capabilities, and deployment trade offs. arXiv preprint arXiv:251003847 10.48550/arXiv.2510.03847

Sima AC, Mendes de Farias T, Zbinden E, et al (2019) Enabling semantic queries across federated bioinformatics databases. Database 2019:baz106. 10.1093/database/baz106

Singh A, Ehtesham A, Kumar S, et al (2025) Agentic retrieval-augmented generation: A survey on agentic rag. arXiv preprint arXiv:250109136 10.48550/arXiv.2501.09136

Slabinoha M, Melnychuk S, Manuliak I, et al (2022) Comparative analysis of embedded databases performance on single board computer systems using python. In: 2022 IEEE 17th International Conference on Computer Sciences and Information Technologies (CSIT), IEEE, pp 222–225

Sloan Z, Arends D, Broman KW, et al (2016) Genenetwork: framework for web-based genetics. Journal of Open Source Software 1(2):25. 10.21105/joss.00025

Smeros P, Emonet V, Wang R, et al (2025) Sparql-LLM: Real-time SPARQL query generation from natural language questions. arXiv preprint arXiv:251214277 10.48550/arXiv.2512.14277

Subramanian S, Akinfaderin A, Zhang Y, et al (2025) Keyword search is all you need: Achieving RAG-Level performance without vector databases using agentic tool use. arXiv preprint arXiv:260223368 10.48550/arXiv.2602.23368

Tomaszuk D, Hyland-Wood D (2020) Rdf 1.1: Knowledge representation and data integration language for the web. Symmetry 12(1):84. 10.3390/sym12010084

Watson PM, Ashbrook DG (2020) Genenetwork: a continuously updated tool for systems genetics analyses. bioRxiv pp 2020–12. 10.1101/2020.12.23.424047

Williams R (1994) The portable dictionary of the mouse genome: a personal database for gene mapping and molecular biology. Mammalian Genome 5(6):372–375. 10.1007/BF00356557

Wysocki O, Carvalho D, Bogatu A, et al (2024) An llm-based knowledge synthesis and scientific reasoning framework for biomedical discovery. In: Proceedings of the 62nd Annual Meeting of the Association for Computational Linguistics (Volume 3: System Demonstrations), pp 355–364, 10.18653/v1/2024.acl-demos.34

Yacouby R, Axman D (2020) Probabilistic extension of precision, recall, and f1 score for more thorough evaluation of classification models. In: Proceedings of the first workshop on evaluation and comparison of NLP systems, pp 79–91, 10.18653/v1/2020.eval4nlp-1.9

Zhavoronkov A, Naumov V, Sidorenko D, et al (2026) An open benchmark and language models for AI in aging biology. Cell 189(19):5980–5994. 10.1016/j.cell.2026.08.026

Zhou J, Li H, Chen S, et al (2025) Large language models in biomedicine and healthcare. npj Artificial Intelligence 1(1):44. 10.1038/s44387-025-00047-1

Zou L, Huang R, Wang H, et al (2014) Natural language question answering over RDF: a graph data driven approach. In: Proceedings of the 2014 ACM SIGMOD international conference on Management of data, pp 313–324, 10.1145/2588555.2610525

